# Causal adaptation to visual input dynamics governs the development of complex cells in V1

**DOI:** 10.1101/756668

**Authors:** Giulio Matteucci, Davide Zoccolan

## Abstract

Visual perception relies on cortical representations of visual objects that remain relatively stable with respect to the variation in object appearance typically encountered during natural vision (e.g., because of position changes). Such stability, known as *transformation tolerance*, is built incrementally along the *ventral stream* (the cortical hierarchy devoted to shape processing), but early evidence of position tolerance is already found in primary visual cortex (V1) for *complex cells*^1^. To date, it remains unknown what mechanisms drive the development of this class of neurons, as well as the emergence of tolerance across the ventral stream. Leading theories suggest that tolerance is learned, in an unsupervised manner, either from the temporal continuity of natural visual experience^2–10^ or from the spatial statistics of natural scenes^11,12^. However, neither learning principle has been empirically proven to be at work in the postnatal developing cortex. Here we show that passive exposure to temporally continuous visual inputs during early postnatal life is essential for normal development of complex cells in rat V1. This was causally demonstrated by rearing newborn rats with frame-scrambled versions of natural movies, resulting in temporally unstructured visual input, but with unaltered, natural spatial statistics. This led to a strong reduction of the fraction of complex cells, which also displayed an abnormally fast response dynamics and a reduced ability to support stable decoding of stimulus orientation over time. Conversely, our manipulation did not prevent the development of *simple cells*, which showed orientation tuning and multi-lobed, Gabor-like receptive fields as sharp as those found in rats reared with temporally continuous natural movies. Overall, these findings causally implicate unsupervised temporal learning in the postnatal development of transformation tolerance but not of shape tuning, in agreement with theories that place the latter under the control of unsupervised adaptation to spatial, rather than temporal, image statistics^13–16^.

---

It has long been proposed that the tuning of sensory neurons is determined by adaptation to the statistics of the signals they need to encode^14,15^. In the visual domain, this notion has given rise to two broad families of unsupervised learning algorithms: those relying on the spatial structure of natural images, referred to as *unsupervised spatial learning* (USL) models^11–16^; and those leveraging the spatiotemporal structure of natural image sequences, referred to as *unsupervised temporal learning* (UTL) models^2–10^. Both kinds of learning have been applied to explain the emergence of shape tuning (or selectivity) and transformation tolerance (or invariance) in the ventral stream – i.e., the ability of ventral neurons to selectively respond to specific visual patterns, while tolerating variations in their appearance^1^. In sparse coding theories (arguably the most popular incarnation of USL), maximizing the sparsity of the representation of natural images produces Gabor-like edge-detectors that closely resemble the receptive fields (RFs) of V1 simple cells^13,16^. Other USL models, by optimizing objective functions that depend on the combination of several linear spatial filters, also account for the emergence of position-tolerant edge detectors, such as V1 complex cells^11,12^. The latter, however, have been more commonly modeled as the result of UTL, where the natural tendency of different object views to occur nearby in time is used to factor out object identity from other faster-varying, lower-level visual attributes. Interestingly, while some UTL models presuppose the existence of a bank of simple cells, upon which the complex cells’ representation is learned^2,6–10^, other models, such as *slow feature analysis* (SFA), directly evolve complex cells from the pixel (i.e., retinal) representation, thus simultaneously learning shape selectivity and invariance^3,4^. To date, it remains unclear what role these hypothesized learning mechanisms play in the developing visual cortex, if any. In fact, although it is well established that early visual experience can strongly affect the development of visual cortical tuning^17–19^, empirical support for the role of sparse coding in determining orientation selectivity is still inconclusive^16,20^, while no causal evidence has been gathered yet to demonstrate the involvement of UTL in postnatal development of invariance and/or selectivity (but see ^21,22^ for a behavioral study of the role of UTL in the development of chicks’ object vision).

To address the latter question, we took 18 newborn rats (housed in light-proof cabinets from birth) and, from postnatal day (P)14 (i.e., at eye opening) through P60 (i.e., well beyond the end of the critical period), we subjected them to daily, 4-hours-long exposures inside controlled visual environments (Fig. 1a). Specifically, 8 animals (the *control* group) were exposed to natural movies, while the remaining 10 (the *experimental* group) were exposed to their frame-scrambled versions (Fig. 1b). Critically, this manipulation destroyed the temporal continuity of natural visual experience for the experimental rats, while sparing the spatial structure of the individual image frames, which remained the same as for the control animals. This allowed isolating the pure contribution of temporal contiguity to the postnatal development of V1 simple and complex cells.

**Fig. 1.**
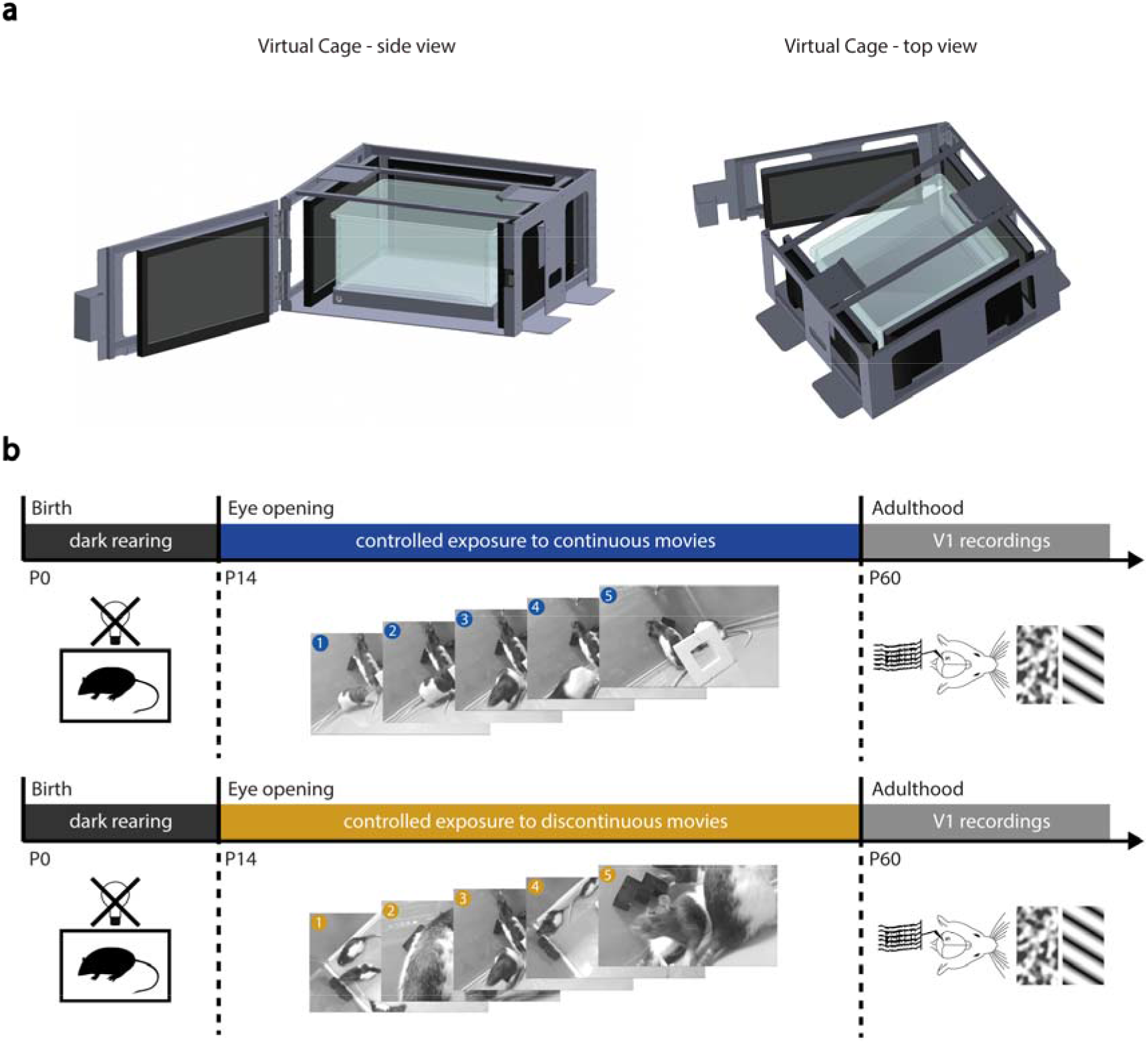
Experimental design. **a**, CAD drawing of a *virtual cage* – the custom apparatus used to rear newborn rats in visually controlled environments. The cage consists of a metal, box-like structure (light gray) holding 4 computer monitors (black/dark gray) that fully surround a transparent basin (light blue), where a newborn rat can be placed for immersive exposure to the visual scenes/movies displayed on the monitors. **b**, Time course of the experiment. Two groups of rats, control (top) and experimental (bottom), were born in dark and housed in lightproof cabinets till eye opening (black bars). Afterward, the control rats were subjected to daily, 4-hours long exposures to natural videos inside the virtual cages (blue bar), while the experimental rats were subjected to the frame-scrambled versions of the same movies (orange bar). Starting from P60, neuronal recordings from V1 of both the control and experimental rats were performed under anesthesia (gray bars), while the animals were exposed to drifting gratings and movies of spatially and temporally correlated noise.

The latter were studied by performing multi-channel extracellular recordings from V1 of each rat under fentanyl/medetomidin anesthesia^23^, shortly after the end of the controlled-rearing period. Our recordings mainly targeted layer 5, which is known to contain the largest fraction of complex cells in rodents^24^ as well as layer 4, with the distributions of recorded units across the cortical depth and the cortical laminae being statistically the same for the control and experimental groups (Extended Data Fig. 1). During a recording session, each animal was presented with drifting gratings spanning 12 directions, and with contrast-modulated movies of spatially and temporally correlated noise^23,24^. Responses to the noise movies allowed inferring the linear RF structure of the recorded units using the spike-triggered average (STA) analysis (see Methods). Responses to the drifting gratings were used to estimate the tuning of the neurons with standard orientation and direction selectivity indexes (OSI and DSI; defined in Fig. 2), as well as to probe their sensitivity to phase-shifts of their preferred gratings, thus measuring their position tolerance^23,24^.

**Fig. 2.**
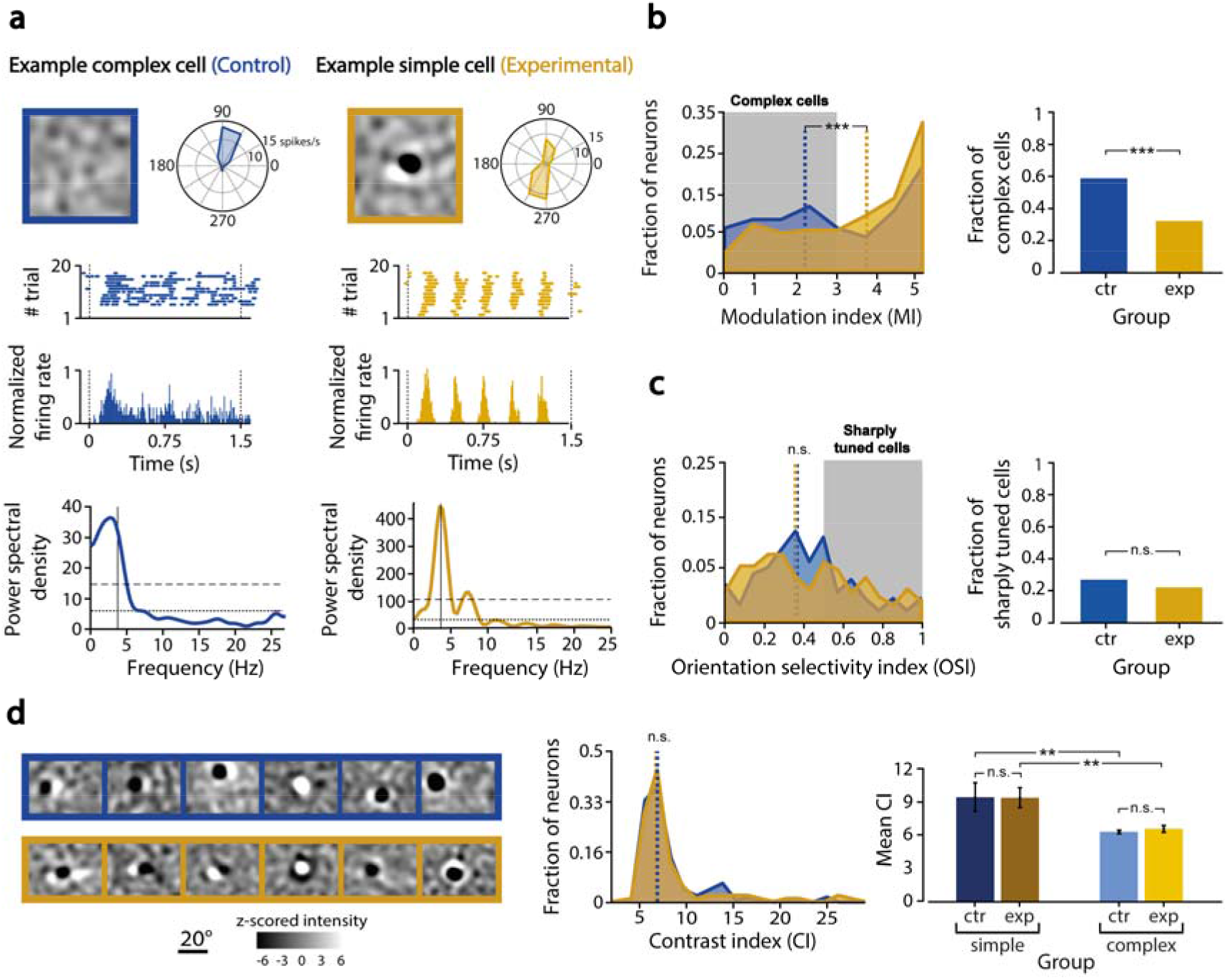
Postnatal rearing in temporally discontinuous visual environments results in an impoverished population of V1 complex cells, but leaves spatial tuning of V1 neurons unaltered. **a**, A representative V1 complex cell of the control group (Left: blue lines) is compared to a representative simple cell of the experimental group (Right: orange lines). For each neuron, the graph shows, from top/left to bottom: 1) the linear RF structure inferred through STA; 2) the direction tuning curve; 3) the raster plot, with the number of spikes (dots) fired across repeated presentations of most effective grating stimulus; 4) the corresponding peri-stimulus time histogram (PSTH), computed in 10 ms-wide time bins; and 5) its power spectrum, with indicated its mean (dotted line), its mean plus SD (dashed line), and the 4 HZ frequency of the grating stimulus (vertical line). **b**, Left: distributions of the modulation index (MI) used to distinguish the poorly phase-modulated complex cells (MI < 3; gray-shaded area) from the strongly modulated simple cells (MI > 3), as obtained for the control (blue) and experimental (orange) V1 populations (only units with OSI > 0.4 included). Both the distributions and their medians (dashed lines) were significantly different (*p* < 0.02, Kolmogorov-Smirnov test; *p* < 0.001, Wilcoxon test). Right: the fraction of units that were classified as complex cells (i.e., with MI < 3) was significantly larger for the control than for the experimental group (*p* < 0.001, Fisher exact-test). **c**, Left: distributions of the orientation selectivity index (OSI), as obtained for the control (blue) and experimental (orange) V1 populations. No significant difference was found between the two distributions and their medians (*p* > 0.05, Kolmogorov-Smirnov test; *p* > 0.05, Wilcoxon test). Right: the fraction of sharply orientation-tuned units (i.e., units with OSI > 0.6) did not differ between the two groups (*p* > 0.05, Fisher exact-test). **d**, Left: Examples of linear RFs inferred through STA for the control (blue frame) and experimental group (orange frame). Center: distributions of the contrast index (CI) used to measure the sharpness of the STA images, as obtained for the control (blue) and experimental (orange) V1 populations. No significant difference was found between the two distributions and their medians (*p* > 0.05, Kolmogorov-Smirnov test; *p* > 0.05, Wilcoxon test). Right: Mean values (± SEM) of the CIs, computed separately for the simple (dark bars) and complex (light bars) cells of the two groups (only units with OSI > 0.4 included). Within each group, the mean CI was significantly larger for the simple than for the complex cells (** *p* < 0.01, two-tailed, unpaired t test).

This is illustrated in Fig. 2a, which shows a representative complex cell from the control group (left: blue lines) and representative simple cell from the experimental group (right: orange lines). Both units displayed sharp orientation tuning (polar plots), but the STA method successfully recovered a sharp, Gabor-like RF only for the simple cell (as expected, given the nonlinear stimulus-response relationship of complex cells^23^). Consistently, the response of the complex cell was only weakly modulated at the temporal frequency (4 Hz) of its preferred grating (middle plots), with the highest power spectral density concentrated at frequencies < 4 Hz (bottom plot). By contrast, the response of the simple cell was strongly phase modulated, with a power spectrum narrowly peaked at the grating frequency. Thus, by z-scoring the power spectral density of the response at the preferred grating frequency, it was possible to define a modulation index (MI) that distinguished between complex (MI < 3) and simple (MI > 3) cells^23^ (see Methods).

We applied this criterion to the neuronal populations of 105 and 158 well-isolated single units recorded from, respectively, the control and experimental group, and we found a significantly lower fraction of complex cells in the latter (39%, 47/158) with respect to the former (55%, 58/105; *p* < 0.01, Fisher exact-test). Consistently, the median MI for the control population (2.69 ± 0.29) was significantly smaller than for the experimental one (3.52 ± 0.25; *p* < 0.05, Wilcoxon test). Such a difference became very sharp after restricting the comparison to the neurons that, in both populations, were at least moderately orientation tuned (i.e., 50 control and 75 experimental units with OSI > 0.4). The resulting MI distribution for the control group had a typical double-peak shape^23^, featuring two maxima, at MI~2 and MI~5, corresponding to the two classes of the complex and simple cells (Fig. 2b, blue curve). Instead, for the experimental group, the peak at low MI was flattened out, leaving a single, prominent peak at MI~5 (orange curve). This resulted in a large, significant difference between the two distributions and their medians (dashed lines), with the fraction of complex cells being almost half in the experimental (35%, orange bar) than in the control group (60%, blue bar). Conversely, no difference was observed between the two groups in terms of orientation tuning (Fig. 2c), with the OSI distributions (blue and orange curves) and their medians (dashed lines) being statistically undistinguishable, as well as the fraction of sharply orientation-tuned units (i.e., neurons with OSI > 0.6; blue vs. orange bar). A similar result was found for direction tuning (Extended Data Fig. 2).

Taken together, these findings suggest that our experimental manipulation substantially impaired the development of complex cells, but not the emergence of orientation and motion sensitivity, and the development of simple cells. This was confirmed by the fact that STA was as successful at yielding sharp, linear RFs (often similar to Gabor filters) for the experimental units as for the control ones (see examples in Fig. 2d, left). The sharpness of the STA images, as assessed through an expressly devised contrast index^23^ (CI; see Methods), was similar for the two groups, with the CI distributions and their medians being statistically undistinguishable (Fig. 2d, blue vs. orange curve/line). As expected, for both groups, the mean CI was significantly larger for the simple than for the complex cells (dark vs. light bars), reflecting the better success of STA at inferring the linear RFs of the former, but no difference was found between the mean CIs of the simple cells of the two groups (dark blue vs. brown bar) and the mean CIs of the complex cells (light blue vs. yellow bar).

Next, we tested the extent to which the experimental units that had been classified as complex cells fully retained the functional properties of this class of neurons. Based on intuitive considerations (i.e., the local invariance of complex cells) and the predictions of SFA^3,4^, complex cells should fire more persistently than simple cells, in response to a continuous, spatiotemporally correlated visual input. To measure the persistence (or slowness) of neuronal responses, we computed the time-constants of the exponential fits to the autocorrelograms of the spike trains evoked by the noise movies (see examples in Fig. 3a). As expected, the average time constant was larger for the control than for the experimental units (Fig. 3b). Such difference, however, was not merely driven by the larger fraction of complex cells in the control group (Fig. 2b). In fact, while no difference of time constant was found between the simple cells of the two groups (Fig. 3c, dark blue vs. brown bar), the firing of complex cells was significantly faster for the experimental than for the control units (yellow vs. light blue bar).

**Fig. 3.**
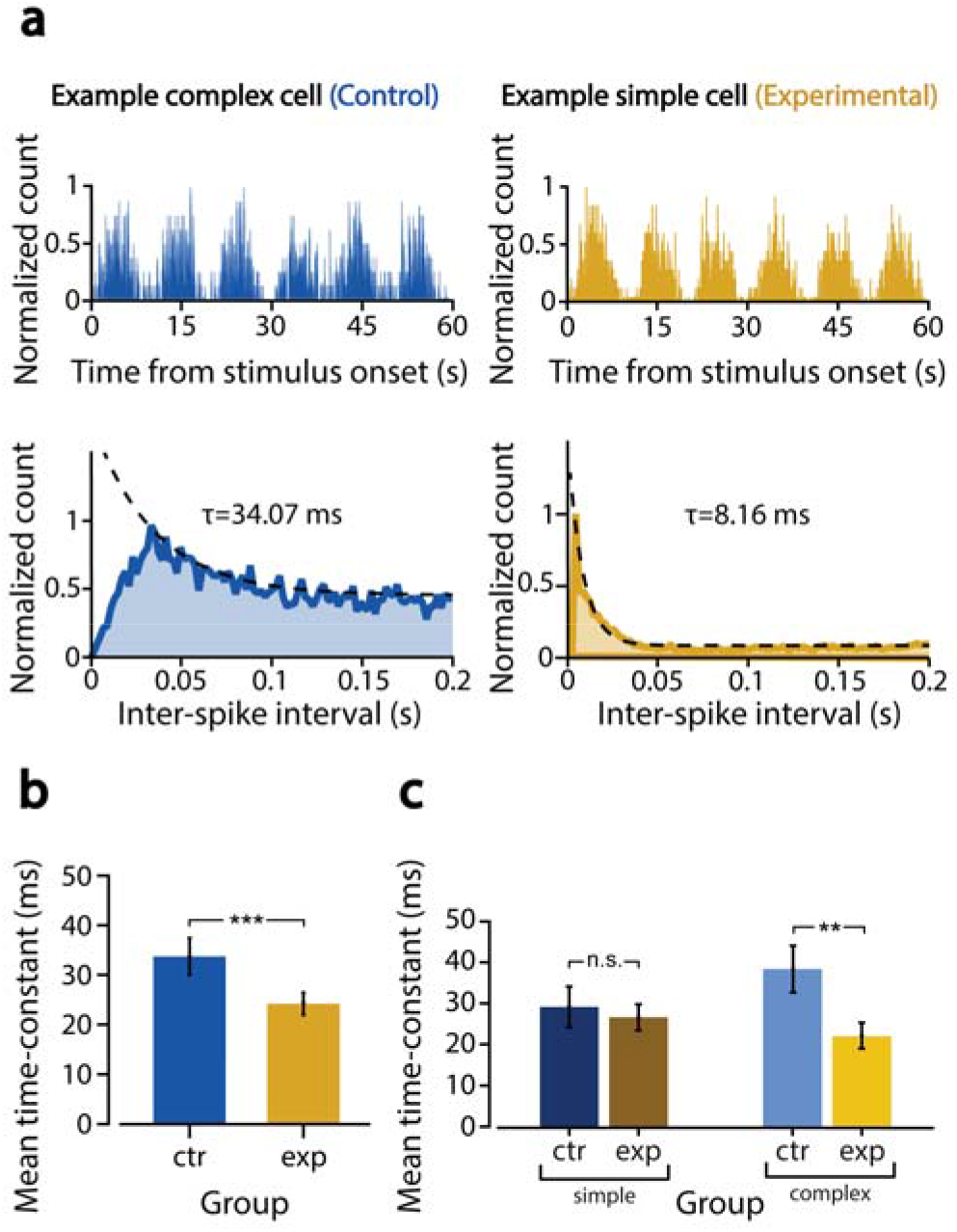
Postnatal rearing in temporally discontinuous visual environments leads to the development of complex cells with abnormally fast response dynamics. **a**, Top: PSTHs showing the average responses of the two example neurons of Fig. 2a to the contrast-modulated noise movies (see Methods). For both neurons, the firing rate was strongly modulated at the frequency of variation of the contrast of the movies (i.e., 0.1 Hz). Bottom: distributions of inter-spike intervals (ISIs) of the spike trains evoked by the noise movies for the two example neurons. The resulting autocorrelograms were fitted with exponential decaying functions (dashed lines) to measure the slowness (i.e., the time constant *τ* of the exponential fit) of the responses. In this example, the complex cell (blue curve) displays slower dynamics (i.e., larger *τ)* than the simple cell (orange curve). The two units also differ in the number of counts at low ISIs, which is much larger for the simple cell, as expected for a unit firing tightly packed trains of spikes (see Fig. 2a). **b**, Mean values (± SEM) of the time constants *τ,* computed for the control (blue) and experimental (orange) populations (*** *p* < 0.001, two-tailed, unpaired t test). **c**, Mean values (± SEM) of the time constants *τ,* computed separately for the simple (dark bars) and complex (light bars) cells of the two groups. While the simple cells had equally fast dynamics, the complex cells were significantly slower in the control than in the experimental group (** *p* < 0.01, two-tailed, unpaired t test).

To understand the functional implication of such abnormally fast dynamics, we assessed the ability of the four distinct populations of simple and complex cells of the two groups to support stable decoding of stimulus orientation over time. To this aim, we randomly sampled 300 neurons from each population (after having first matched the populations in terms of OSI and orientation preference distributions; see Methods), so as to obtain four, equally sized and similarly tuned pseudo-populations, whose units homogenously covered the orientation axis. We then trained binary logistic classifiers to discriminate between 0°- and 90°-oriented gratings (drifting at 4 Hz), based on the activity of each pseudopopulation. Each classifier was trained using neuronal responses (spike counts) in a 33 ms-wide time bin that was randomly chosen within the presentation epoch of the gratings. We then tested the ability of each classifier to generalize the discrimination to test bins at increasingly larger time lags (TLs) from the training bin (see Fig. 4a and Methods for details). As expected, given the strong phase dependence of their responses (see cartoon in Fig. 4a, top), the simple cells from both groups yielded generalization curves that were strongly modulated over time and virtually identical (Fig. 4b, dark blue and brown curves). The performance was high (≥ 80% correct) at test bins where the phase of the grating was close to that of the training bin (i.e., at TLs that were multiple of the 250 ms grating period), but it dropped to less than 30% correct (i.e., well below chance; dashed line) at test bins where the grating was in opposition of phase with respect to the training bin (e.g., at TL ~ 125 ms). By comparison, the complex cells of the control group, by virtue of their weaker phase dependence (see cartoon in Fig. 4a, bottom), afforded a way more phase-tolerant decoding of grating orientation, with the performance curve never dropping below chance level at any TL (Fig. 4b, light blue curve). However, for the complex cells of the experimental group, the performance curve (in yellow) was not as stable – at most TLs, it was 5-10 percentage points smaller than the performance yielded by the control complex cells,dropping significantly below chance at test bins where the grating was in opposition of phase with respect to the training bin. That is, the ability of the experimental complex cells to support phase-tolerant orientation decoding was somewhat in between that of properly developed complex cells and that of simple cells.

**Fig. 4.**
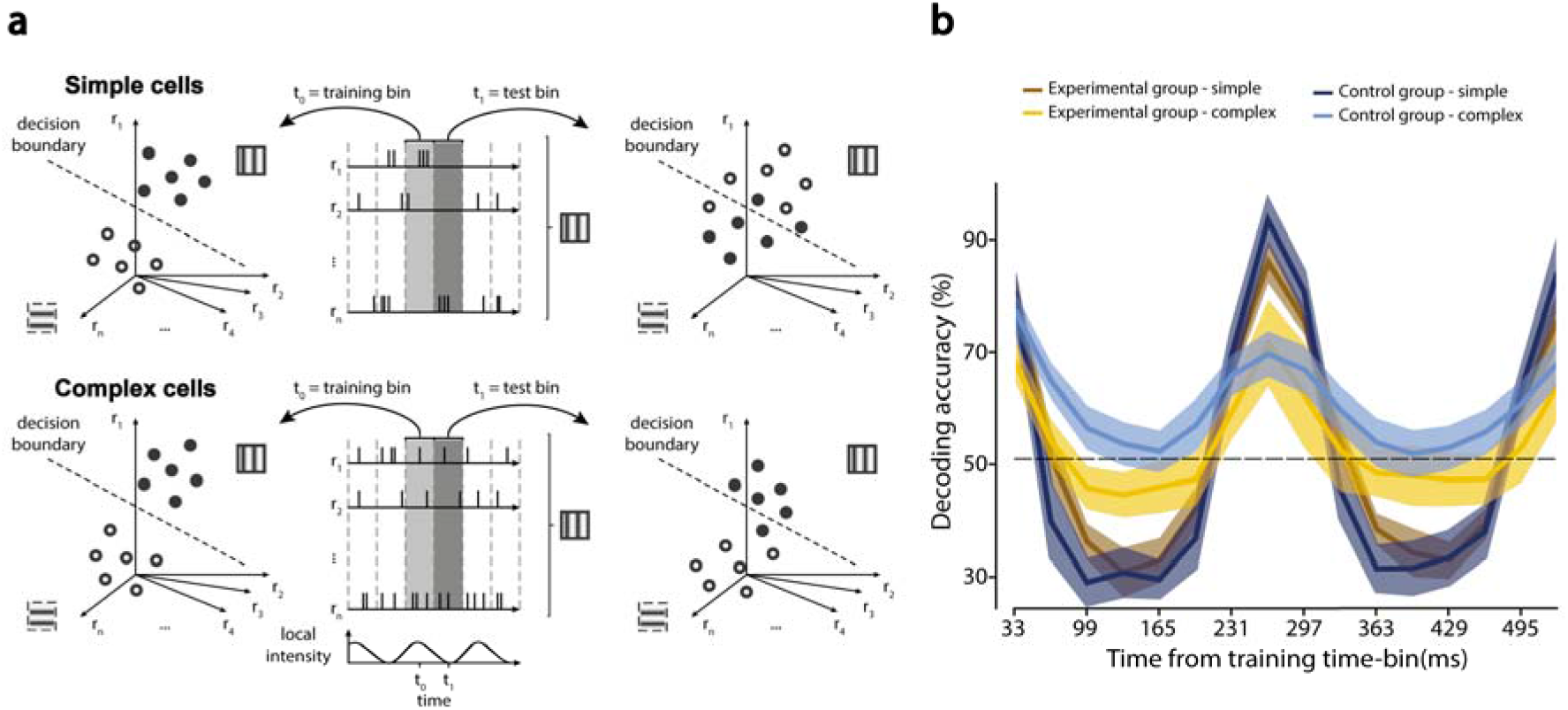
Postnatal rearing in temporally discontinuous visual environments reduces the ability of complex cells to support phase-tolerant discrimination of grating orientation. **a**, The cartoon illustrates the expected outcome of the decoding analysis to test the ability of simple and complex cells to support phase-tolerant discrimination of grating orientation. In the case of simple cells (top), a linear classifier built at time *t*_0_ (middle: light gray shading) to successfully discriminate a vertical from a horizontal drifting grating (left: the filled and empty dots are well separated, within the neuronal representational space, by the linear decision boundary) will generalize poorly, when tested at a later time *t*_1_ (middle: dark gray shading), with the accuracy dropping even below chance (right: the filled and empty dots swap sides of the linear decision boundary), because of the strong phase dependency of the responses *r_i_* (middle: some neurons firing at *t*_0_ will stop firing at *t*_1_, while some other units that are silent at *t*_0_ will respond at *t*_1_). By contrast, for a population of complex cells (bottom), given the larger stability of the responses *r_i_* (middle), the decision boundary resulting from the training at *t*_0_ (left) will generalize better at *t*_1_ (right: the filled and empty dots are still mostly on the original side of the decision boundary). **b**, Decoding accuracy yielded by the four populations of simple control (dark blue), complex control (light blue), simple experimental (brown) and complex experimental (orange) cells, in the vertical (i.e., 90°) vs. horizontal (i.e., 0°) grating discrimination task, in 33 ms-wide test bins located at increasingly larger time lags from the training bin (i.e., bin with lag = 0). The solid curves are the averages of many resampling loops of the neuronal population vectors and the training bins (see Methods). The shaded regions are the bootstrap-estimated 95% confidence intervals of the solid lines (see Methods).

Overall, these findings show that destroying the temporal continuity of early visual experience severely interferes with the typical development of complex cells in V1, leading to a sizable reduction of their number (Fig. 2b) and an impairment of their functional properties (Fig. 3c and 4b). This implies that experience with the temporal structure of natural image sequences plays a critical role in the postnatal development of the earliest form of invariance found along the ventral stream – a role that, so far, had been empirically demonstrated only in adult monkeys, at the very last stage of this pathway: the inferotemporal cortex^25,26^. At the same time, our experiments show that development of orientating tuning is unaffected by the lack of experience with temporal continuity (Fig. 2d), with simple cells exhibiting unaltered spatial (Fig. 2c), temporal (Fig. 3c) and functional (Fig. 4b) properties.

From a theoretical standpoint, these results causally validate the family of UTL models^2–10^ at the neural level, albeit limiting their scope to the development of invariance. More in general, since slowness has been related to predictability^27,28^, our results are also consistent with normative approaches to sensory processing that are based on temporal prediction^29^. On the other hand, our findings, by showing that exposure to the spatial structure of natural images alone is not enough to yield complex cells, reject computational accounts of invariance based on USL^11,12^, while leaving open the possibility that the latter may govern the development of shape tuning^13–16^. As a result, our study tightly constrains unsupervised models of visual cortical development, supporting theoretical frameworks where the objectives of sparseness and slowness maximization coexist to yield, respectively, shape selectivity and transformation tolerance ^8,9,30^.

## Methods

All animal procedures were in agreement with international and institutional standards for the care and use of animals in research and were approved by the Italian Ministry of Health (Project DGSAF 22791-A, submitted on September 7, 2015 and approved on December 10, 2015, approval 1254/2015-PR).

### Animal subjects and controlled rearing protocol

Data were obtained from 18 Long–Evans male rats that were born and reared in our facility for visually controlled rearing. The facility consists of a small vestibule, where the investigators can wear the infrared goggles that are necessary to operate in total darkness, and a larger, lightproof room, containing a lightproof housing cabinet (Tecniplast) and four custom cabinets (Tecniplast) for exposure of the rats to controlled visual environments.

Pregnant mothers (Charles River Laboratories) where brought into the housing cabinet about one week before delivery. Pups were born inside the cabinet and spent the first two weeks of their life in total darkness with their mothers. Starting from P14 (i.e., at eye opening) through P60 (i.e., well beyond the end of the critical period), each rat, while still housed in full darkness (i.e., inside the housing cabinet) for most of the day, was also subjected to daily, 4-hours-long exposures inside an immersive visual environment (referred to as the *virtual cage*), consisting of a transparent basin (480×365×210 mm, Tecniplast 1500U), fully surrounded by 4 computer-controlled LCD monitors (one per wall; 20” HP P202va; see Fig. 1A), placed on the shelf of one of the custom cabinets (each cabinet had 4 shelves, for a total of 16 rats that could be simultaneously placed in the visually controlled environments). These controlled-rearing environments (which are reminiscent of those used to study the development of object vision in chicks^21,22,31^) were custom designed in collaboration with Videosystem, which took care of building and installing them inside the custom cabinets.

Different visual stimuli were played on the monitors, depending on whether an animal was assigned to the experimental or the control group. Rats in the control group (*n* = 8) were exposed to natural movies, including both indoor and outdoor scenes, camera self-motion and moving objects. Overall, the rearing playlist included 16 videos of different duration, lasting from a few minutes to half an hour. The playlist was played in random order and looped for the whole duration of the exposure. Rats from the experimental group (*n* = 10) were exposed to a time-shuffled version of the same movies, where the order of the frames within each video was randomly permuted, so as to destroy the temporal continuity of the movie, while leaving unaltered the natural spatial statistics of the individual image frames.

All movies were played at 15 Hz, which is approximately half of the critical flicker fusion frequency (~30-40 Hz) that has been measured for the rat^32^. This ensured that, while the temporal correlation of the input was substantially broken, no fusion occurred between consecutive frames of the movies, thus allowing the rats to experience the spatial content of the individual image frames. On the other hand, choosing a frame rate lower than the flicker fusion frequency allowed the rats, at least in principle, to still experience some residual amount of temporal continuity in the visual input, resulting from scanning the image frames trough head or eye movements. This, together with the presence of some stable visual features in the environment (e.g., the dark edges of the monitors) and the possibility for the rats to see their own body, may account for the residual fraction of complex cells observed in experimental group (Fig. 2B).

Animal care, handling and transfer operations were always executed in absolute darkness, using night vision goggles (Armasight NXY7), in such a way to prevent any unwanted exposure of the animals to visual inputs different from those chosen for the rearing.

### Surgery and recordings

Acute extracellular recordings were performed between P60 and P90 (last recording). During this 30-days period, the animals waiting to undergo the recording procedure were maintained on a reduced visual exposure regime (i.e., 2-hours-long visual exposure sessions every second day; see previous section).

The surgery and recording procedure was the same as described in ^23^. Briefly, the day of the experiment, the rat was taken from the rearing facility and immediately (within 5-10 minutes) anesthetized with an intraperitoneal injection of a solution of 0.3 mg/kg of fentanyl (Fentanest, Pfizer) and 0.3 mg/kg of medetomidin (Domitor, Orion Pharma). A constant level of anesthesia was then maintained through continuous intraperitoneal infusion of the same aesthetic solution used for induction, but at a lower concentration (0.1 mg/kg/h fentanyl and 0.1 g/kg/h medetomidine), by means of a syringe pump (NE-1000; New Era Pump Systems). After induction, the rat was secured to a stereotaxic apparatus (Narishige, SR-5R) in flat-skull orientation (i.e., with the surface of the skull parallel to the base of the stereotax) and, following a scalp incision, a craniotomy was performed over the target area in the left hemisphere (typically, a 2×2 mm^2^ window), and the dura was removed to allow the insertion of the electrode array. The coordinates of penetration used to target V1 were ~6.5 mm posterior from bregma and ~4.5 mm left to the sagittal suture (i.e., anteroposterior 6.5, mediolateral 4.5). Once the surgical procedure was completed, and before probe insertion, the stereotax was placed on a rotating platform and the rat’s left eye was covered with black, opaque tape, while the right eye (placed at 30 cm distance from the monitor) was immobilized using a metal eye-ring anchored to the stereotax. The platform was then rotated in such a way to bring the binocular visual field of the right eye to cover the left side of the display.

Extracellular recordings were performed using either single- (or double-) shank 32- (or 64-) channel silicon probes (NeuroNexus Technologies) with site recording area of 775 μm^2^ and 25 μm of intersite spacing. After grounding (by wiring the probe to the animal’s head skin), the electrode was manually lowered into the cortical tissue using an oil hydraulic micromanipulator (Narishige, MO-10; typical insertion speed: 5 μm/s), up to the chosen insertion depth (800 –1000 m from the cortical surface), either perpendicularly or with a variable tilt, between 10° and 30°, relative to the vertical to the surface of the skull. Extracellular signals were acquired using a system 3 workstation (Tucker Davis Technologies) with a sampling rate of 25 kHz.

Since, in rodents, the largest fraction of complex cells in found in layer 5 of V1^24^, our recordings aimed at sampling more densely that layer. This was verified *a posteriori* (Extended Data Fig. 1), by estimating the cortical depth and laminar location of the recorded units, based on the patterns of *visually evoked potentials* (VEPs) recorded across the silicon probes used in our recording sessions. More specifically, we used a template-matching algorithm for laminar identification of cortical recording sites that we recently developed and validated in an appositely dedicated methodological study^33^. Briefly, the method finds the optimal match between the pattern of VEPs recorded, in a given experiment, across a silicon probe and a template VEP profile, spanning the whole cortical thickness, that had been computed by merging an independent pool of 18 recording sessions, in which the ground-true depth and laminar location of the recording sites had been recovered through histology. The method achieves a cross-validated accuracy of 79 μm in recovering the cortical depth of the recording sites and a 72% accuracy in returning their laminar position, with the latter increasing to 83% for a coarser grouping of the layers into supagranular (1-3), granular (4) and infragranular (5-6).

### Visual stimuli

During a recording session, each animal was presented with: i) 20 repetitions (trials) of 1.5-s-long drifting gratings, made of all possible combinations of two spatial frequencies (0.02 and 0.04 cpd), two temporal frequencies (2 and 4 Hz), and 12 directions (from 0° to 330°, in 30° increments); and ii) 20 different 60-s-long spatially and temporally correlated, contrast modulated, noise movies^23,24^. All stimuli were randomly interleaved, with a 1-s-long interstimulus interval (ISI), during which the display was set to a uniform, middle-gray luminance level. To generate the movies, random white noise movies were spatially correlated by convolving them with a Gaussian kernel having full width at half maximum (FWHM) corresponding to a spatial frequency of 0.04 cpd SF. Temporal correlation was achieved by convolving the movies with a causal exponential kernel with a 33 ms decay time-constant. To prevent adaptation, each movie was also contrast modulated using a rectified sine wave with a 10 s period from full contrast to full contrast^24^.

Stimuli were generated and controlled in MATLAB (MathWorks) using the Psychophysics Toolbox package and displayed with gamma correction on a 47-inch LCD monitor (SHARP PNE471R) with 1920×1080-pixel resolution, 220 cd/m^2^ maximum brightness, and spanning a visual angle of 110° azimuth and 60° elevation. Grating stimuli were presented at 60 Hz refresh rate, whereas noise movies were played at 30 Hz.

### Single unit isolation

Single units were isolated offline using the spike sorting package KlustaKwik-Phy^34^. Automated spike detection, feature extraction, and expectation maximization clustering were followed by manual refinement of the sorting using a customized version of the Phy interface. Specifically, we took into consideration many features of the candidate clusters: a) the distance between their centroids and their compactness in the space of the principal components of the waveforms (a key measure of goodness of spike isolation); b) the shape of the auto- and cross correlograms (important to decide whether to merge two clusters or not); c) the variation, over time, of the principal component coefficients of the waveform (important to detect and take into account possible electrode drifts); and d) the shape of the average waveform (to exclude, as artifacts, clearly non physiological signals). Clusters suspected to contain a mixture of one or more single units were separated using the “reclustering” feature of the GUI. After the manual refinement step, we included in our analyses only units that were: i) well-isolated, i.e. with less than 0.5% of “rogue” spikes within 2 ms in their autocorrelogram; and ii) grating-responsive, i.e., with the response to the most effective grating condition being larger than 2 spikes per second (baseline-subtracted), and being larger than 6 z-scored points relative to baseline activity. The average baseline (spontaneous) firing-rate of each well-isolated unit was computed by averaging its spiking activity over every ISI condition. These criteria led to the selection of 105 units for the control group and 158 units for experimental group.

### Quantification of selectivity

The response of a neuron to a given drifting grating was computed by counting the number of spikes during the whole duration of the stimulus, averaging across trials and then subtracting the spontaneous firing rate (see previous section). To quantify the tuning of a neuron for the orientation and direction of drifting gratings, we computed two standard metrics, the orientation and direction selectivity indexes (OSI and DSI), which are defined as: OSI = (*R*_pref_ – *R*_ortho_)/(*R*_pref_), and DSI = (*R*_pref_ – *R*_opposite_)/(*R*_pref_), where *R*_pref_ is the response of the neuron to the preferred direction, *R*_ortho_ is the response to the orthogonal direction, relative to the preferred one (i.e., *R*_ortho_ = *R*_pref_ + *π*/2), and *R*_opposite_ is the response to the opposite direction, relative to the preferred one (i.e., *R*_opposite_ = *R*_pref_ + *π*). Values close to one indicate very sharp tuning, whereas values close to zero are typical of untuned units.

### Quantification of phase modulation (or position tolerance)

Since phase shifts of a grating are equivalent to positional shifts of the whole, 2D sinusoidal pattern, a classical way to assess position tolerance of V1 neurons (thus discriminating between simple and complex cells) is to probe the phase sensitivity of their responses to optimally oriented gratings. Quantitatively, the phase-dependent modulation of the spiking response at the temporal frequency *f*_1_ of a drifting grating was quantified by a modulation index (MI) adapted from^35^ and used in^23^, defined as:

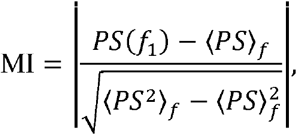

where *PS* indicates the power spectral density of the stimulus-evoked response, i.e., of the peri-stimulus time histogram (PSTH), and 〈 〉_*f*_ denotes the average over frequencies. This metric measures the difference between the power of the response at the stimulus frequency and the average value of the power spectrum in units of its standard deviation. The power spectrum was computed by applying the Blackman-Tukey estimation method to the baseline-subtracted, 10ms-binned PSTH. Being MI a standardized measure, values greater than 3 can be interpreted as signaling a strong modulation of the firing rate at the stimulus frequency (typical of simple cells), whereas values smaller than 3 indicate poor modulation (typical of complex cells). On this ground, we adopted MI = 3 as a threshold for classifying neurons as simple or complex.

### Linear receptive field estimation

We used the spike-triggered average (STA) method^36,37^ to estimate the linear RF structure of each recorded neuron. The method was applied to the spike trains fired by neurons in response to the spatiotemporally correlated and contrast modulated noise movies described above. To account for the correlation structure of our stimulus ensemble and prevent artifactual blurring of the reconstructed filters, we “decorrelated” the raw STA images by dividing them by the covariance matrix of the whole stimulus ensemble^36,37^. We used Tikhonov regularization to handle covariance matrix inversion. Statistical significance of the STA images was then assessed pixel-wise, by applying the following permutation test. After randomly reshuffling the spike times, the STA analysis was repeated multiple times (*n* = 50) to derive a null distribution of intensity values for the case of no linear stimulus-spike relationship. This allowed z-scoring the actual STA intensity values using the mean and standard deviation of this null distribution. The temporal span of the spatiotemporal linear kernel we reconstructed via STA extended till 330 ms before spike generation (corresponding to 10 frames of noise at 30 Hz frame rate). The STA analysis was performed on downsampled noise frames (16×32 pixels) and the resulting filters were later spline-interpolated at higher resolution for better visualization.

To estimate the amount of signal contained in a given STA image, we used the contrast index (CI) metric that we have introduced in a previous study^23^. The CI is a robust measure of maximal local contrast in a z-scored STA image. Since the intensity values of the original STA images were expressed as z-scores (see above), a given CI value can be interpreted in terms of peak-to-peak (i.e. white-to-black) distance in sigma units of the z-scored STA values. For the analysis shown in Fig. 2C, the STA image with the highest CI value was selected for each neuron.

### Quantification of response slowness

For each neuron, we quantified the slowness of its response to the same noise movies used to estimate its RF, by computing the time constant of the autocorrelogram of the evoked spike trains (i.e., the probability density function of inter-spike intervals). Being the noise movies composed of richer visual patterns than drifting gratings (i.e., richer orientation and spatial frequency content), this was a way to assess the response properties of the recorded population in a slightly more naturalistic stimulation regime. The time constants τ were computed by fitting the autocorrelogram with the following exponential function:

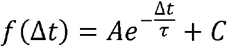

where Δ*t* is the inter-spike time interval (see Fig. 3a, bottom).

Only neurons that were strongly modulated at the frequency of variation of the contrast in the movies were included in this analysis. The level of modulation was quantified by an index similar to the one used to assess the phase sensitivity of responses to the gratings. To this aim, we built PSTHs for the noise movies, by considering each of the 20 different movies we presented as a different trial of the same stimulus, so as to highlight the effect of contrast modulation (see examples of highly contrast modulated neurons in Fig. 3a, top).

### Orientation decoding analysis

The goal of this analysis was to build four pseudo-populations of neurons – i.e., control simple (CS), control complex (CC), experimental simple (ES) and experimental complex (EC) cells – with similar distributions of orientation tuning and orientation preference, and then compare their ability to support stable decoding of the orientation of the gratings over time.

The pseudo-populations were built as follows. We first matched the control and experimental populations in terms of the sharpness of their orientation tuning. To this aim, we took the OSI distributions of the two populations (i.e., the blue and orange curves in Fig. 2c) and, for each bin *b* in which the OSI axis had been divided (i.e., 10 equi-spaced bins of size = 0.1), we took as a reference the population with the lowest number of units *N_b_* in that bin. For this population, all the *N_b_* units were considered, while, for the other population, *N_b_* units were randomly sampled (without replacement) from those with OSI falling in the bin *b*. Repeating this procedure for all the 10 bins, we obtained two down-sampled populations of control and experimental units, having all the same OSI distribution and the same number of units (*n* = 92). When considering separately the pools of simple and complex cells within such down-sampled populations, the resulting mean OSI were very similar (CS: 0.44 ± 0.04, *n* = 43; CC: 0.42 ± 0.03, *n* = 49; ES: 0.46 ± 0.03, *n* = 57; EC: 0.38 ± 0.04, *n* = 35) and not statistically different pairwise (two-tailed, unpaired t-test *p* > 0.05). Matching the four populations in terms of the OSI was essential, but not sufficient, to make sure that they had approximately the same power to support discrimination of the oriented gratings. In fact, the populations could still differ in terms of the distributions of orientation preference. To also equate them in this sense, and make sure that all possible orientations were equally discriminable, we replicated each unit 11 times, by circularly shifting its tuning curve of 11 incremental steps of 30°. This yielded four final pseudo-populations of 473 (CS), 539 (CC), 627 (ES) and 385 (EC) units, with matched orientation tuning and homogeneous orientation preference, to be used for the decoding analysis.

The latter worked as follows. From each pseudo-population, we sampled (without replacement) 300 units (referred to as *decoding pool* in what follows) and we built 300dimensional population vectors having as components the responses (i.e., spike counts) of the sampled units in randomly selected presentations (i.e., trials) of either the 0°- or the 90°-oriented grating (drifting at 4 Hz), with each response computed in the same, randomly chosen 33 ms-wide time bin within the presentation epoch of the grating. More specifically, this time bin was chosen under the constraint of being between 561 ms and 957 ms from the onset of stimulus presentation, so that the drifting grating continued for at least 2 full cycles (i.e., 561 ms) after the selected bin. The random sampling of the trial to be used in a given population vector was performed independently for each neuron (and without replacement), so as to get rid of any noise correlation among the units that were recorded in the same session. Given that 20 repeated trials were recorded per neuron and stimulus condition, a set of 20 population vectors was built for the 0°-oriented grating and another set for the 90°-oriented gratings. These vectors were used to train a binary logistic classifier to discriminate the two stimuli. The resulting classifier was then tested for its ability to discriminate the gratings in 33 ms-wide test bins that were increasingly distant (in time) from the training bin, covering two full cycles of the drifting gratings (i.e., from 33 to 561 ms following the training bin; see abscissa in Fig. 4b). This analysis was repeated for 50 random samplings (without replacement) of the decoding pools and, given a decoding pool, for 10 independent random draws (without replacement) of the training time bin. The resulting 500 accuracy curves were then averaged to yield the final estimate of the stability of the classification over time (solid curves in Fig. 4b).

To obtain 95% confidence intervals (shaded regions in Fig. 4b) for such average classification curves, we run a bootstrap analysis that worked as follows. For each of the four pseudo-populations, we sampled (with replacement) 50 surrogate populations and we used those to re-run the whole decoding analysis described in the previous paragraph. This yielded 50 bootstrap classification curves that were used to compute standard errors for the actual generalization curve. The standard errors were then converted into confidence intervals by multiplying them by the appropriate critical value 1.96 (as described in ^38^).

## Acknowledgments

We thank Mattia D’Andola, Nelly Redolfi and Margherita Riggi for their help with the procedures to rear the rats in the visually controlled environments. We also thank Mattia D’Andola for his support during data collection. We thank Daniele Bertolini for his support in designing and building the apparatus for visually controlled rearing. We thank Laurenz Wiskott for his comments on the manuscript. This work was supported by a European Research Council Consolidator Grant (DZ, project n. 616803-LEARN2SEE).

## Author contributions

GM and DZ designed the study. GM performed and analyzed the neuronal recordings. GM wrote the code for the analyses. GM and DZ wrote the manuscript. GM and DZ revised the manuscript.

## Author information

The authors declare no competing interests.

## Data and code availability

The code to perform the analysis described in the paper as well as the neural data collected for this study will be made available upon request via email to the corresponding author.

## Figures and figure legends

**Extended Data Fig. 1.**
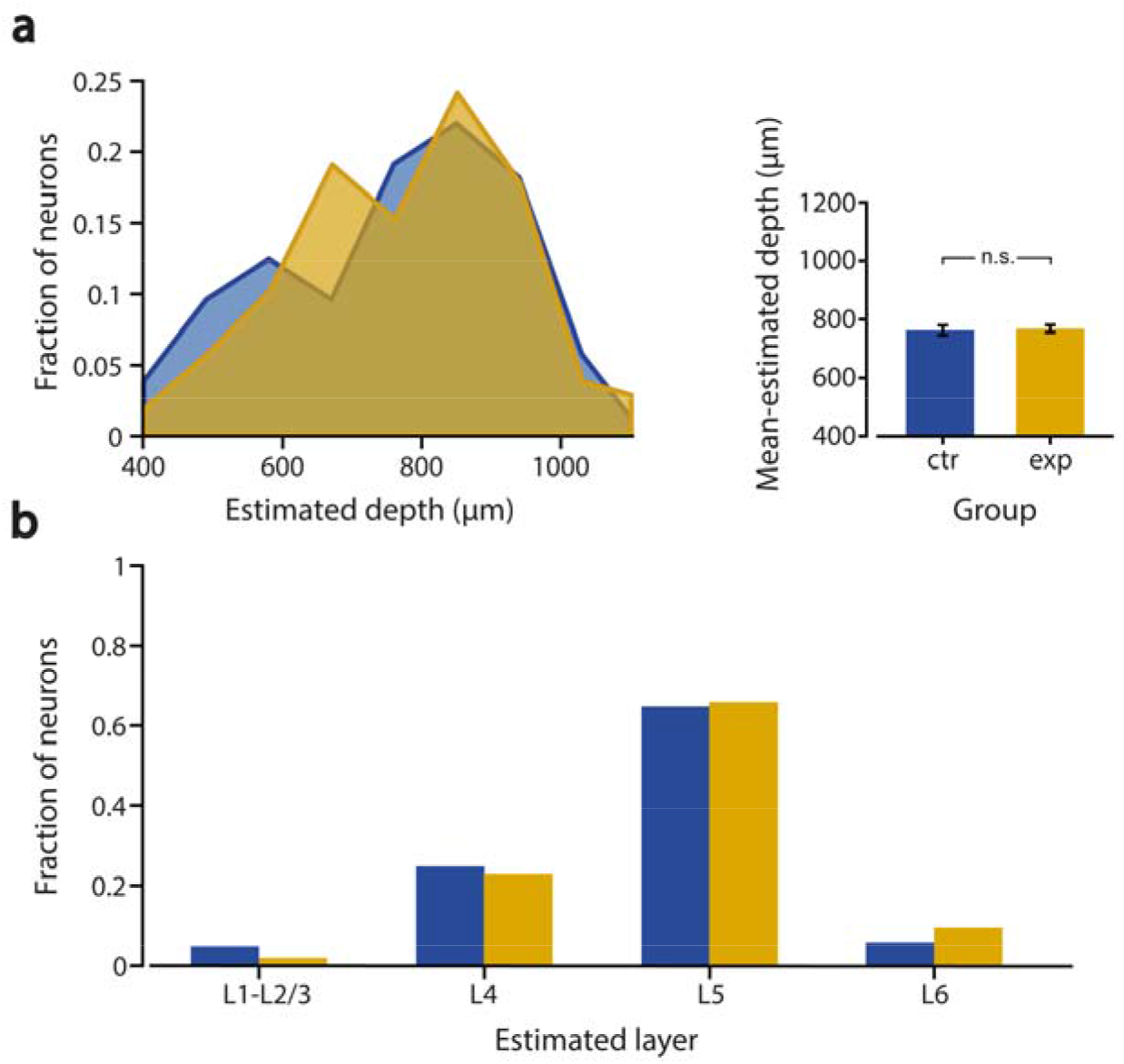
The distributions of cortical depths and laminar locations did not differ between the populations of control and experimental units. **a**, The distributions of cortical depths (left) and their means (right) did not significantly differ between the populations of control (blue) and experimental (orange) units *(p* > 0.05, Kolmogorov-Smirnov test; *p* > 0.05, two-tailed, unpaired t-test). Error bars are SEMs. **c**, The distributions of laminar locations were also not significantly different between control (blue) and experimental (orange) units *(p* > 0.05, χ^2^ test). Both the cortical depths and laminar locations were estimated from the patterns of visually evoked potentials recorded along the silicon probes (see Methods).

**Extended Data Fig. 2.**
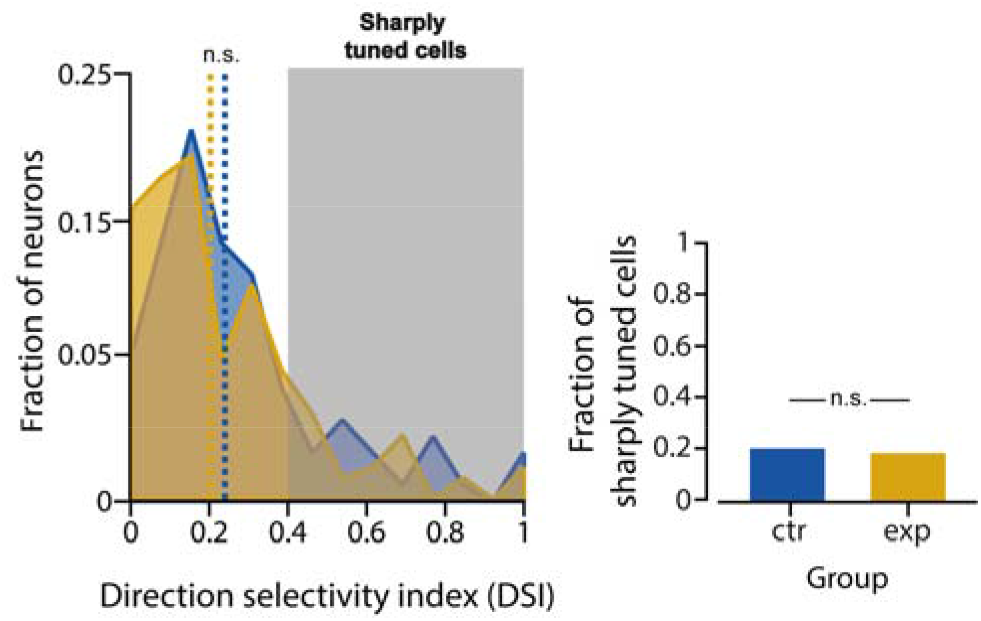
Postnatal rearing in temporally discontinuous visual environments leaves direction tuning of V1 neurons unaltered. Left: distributions of the direction selectivity index (DSI) used to measure the sharpness of orientation tuning, as obtained for the control (blue) and experimental (orange) V1 populations. No significant difference was found between the two distributions and their medians (*p* > 0.05, Kolmogorov-Smirnov test; *p* > 0.05, Wilcoxon test). Right: the fraction of sharply orientation-tuned units (i.e., units with DSI > 0.6) did not differ between the two groups (*p* > 0.05, Fisher exact-test).

